# Imprint of parity and age at first birth on the genomic landscape of subsequent breast cancer

**DOI:** 10.1101/351205

**Authors:** Bastien Nguyen, David Venet, Matteo Lambertini, Christine Desmedt, Roberto Salgado, Hugo Horlings, Françoise Rothé, Christos Sotiriou

## Abstract

**Background:** Although parity and age at first birth are among the most known extrinsic factors that modulates breast cancer risk, their impact on the biology of subsequent breast cancer has never been explored in depth. In this study, we investigate the imprint of parity and age at first birth on the pattern of somatic mutations, somatic copy number alterations (SCNAs), transcriptomic profiles, and tumor infiltrating lymphocytes (TILs) levels of subsequent breast cancer.

**Methods:** A total of 313 patients with primary breast cancer with available whole genome and RNA sequencing data were included in this study. We used a multivariate analysis adjusted for age at diagnosis, pathological stage, molecular subtypes and histological subtypes. We compared nulliparous vs. parous, late parous vs. early parous, and nulliparous vs. pregnancy associated breast cancer (PABC) patients. Late and early parous patients were grouped by using the median age at first birth as a cut-off value. PABC was defined as patients diagnosed up to 10 years postpartum.

**Results:** Genomic alterations of breast cancer are associated with age at first birth but not parity status alone. Independently of clinicopathological features, early parous patients developed tumors characterized by a higher number of Indels (*P*_*adj*_ = 0.002), a lower frequency of *CDH1* mutations (1.2% vs. 12.7% *P*_*adj*_ = 0.013), a higher frequency of *TP53* mutations (50% vs. 22.5%; *P*_*adj*_ = 0.010) and *MYC* amplification (28% vs. 7% *P*_*adj*_ = 0.008), and a lower prevalence of mutational signature 2. PABC were associated with increased TILs infiltration (*P*_*adj*_ = 0.0495).

**Conclusions:** These findings highlight an unprecedented link between reproductive history and the genomic landscape of subsequent breast cancer. With the rapid development of precision oncology, this work advocates that reproductive history should not be underestimated in future clinical studies of breast cancer.

## Background

The effect of parity and age at first birth on the risk of developing breast cancer has been well documented (1–5). Parity is known to have a dual effect on breast cancer risk with an increased risk during 5 to 10 years after pregnancy, followed by a strong and life-long protective effect (1,6). This effect is strongly influenced by age at first birth as pregnancy-induced tumor protection is more pronounced if first birth has occurred early in life. Recent data suggest that pregnancy-induced tumor protection is different according to breast cancer subtypes, with parity and young age at first birth being associated with a marked reduction in the risk of developing luminal subtype tumors (7–10).

Several studies have attempted to investigate the mechanisms underlying this phenomenon (11,12). However, although parity and age at first birth are among the most known extrinsic factors that modulates breast cancer risk, their impact on the biology of breast cancer has never been explored in depth. In the present study, we used a multivariate analysis to investigate the imprint of parity and age at first birth on the pattern of somatic mutations, somatic copy number alterations (SCNAs), transcriptomic profiles, and tumor infiltrating lymphocytes (TILs) levels in a series of 313 breast cancer patients with available whole genome and RNA sequencing data.

## Results

From a publicly available dataset comprising 560 breast cancer patients (13), a total of 313 with available information on parity were included. We identified 264 (84.3%) parous and 49 (15.7%) nulliparous patients (Supplementary Figure S1). In the parous group, 153 patients (57.9%) had available information on age at first birth (median of 25 years, range 16-46 years). Parous patients were divided into two groups: 82 early and 71 late parous patients by using the median age at first birth as a cut-off value. All patients had available somatic mutations and SCNAs data, 182 patients (58.1%) had available transcriptomic data and 170 patients (54.3%) had information on TILs levels (Supplementary Figure S2).

Table 1 summarizes the clinicopathological features of patients. Compared to parous patients, nulliparous patients had significantly larger tumors (tumor size > 2cm, 59.2% vs. 37.1 %, *P* = 0.006), higher frequency of node involvement (40.8% vs. 27.7%; *P* = 0.027) and lower frequency of triple negative disease (TNBC) (4.1 % vs. 23.2%; *P* = 0.001). Compared to early parous patients, late parous patients had a younger age at breast cancer diagnosis (median, 49 years; range, 28-81 years vs. median, 59 years; range, 34-81 years; *P* = 2.58 × 10^−5^) and were more often pre-menopausal (45.3% vs. 20.9%; *P* = 0.005). Late parous patients also had a lower frequency of large tumors (tumor pathological size > 2cm, 40.8% vs. 51.2%, *P* = 0.012), lower frequency of TNBC (19.7% vs. 39%, *P* = 0.026) and higher frequency of lobular histological subtype (14.5% vs. 2.5%, *P* = 0.01). In parous patients, we found a negative correlation between age at first birth and age at breast cancer diagnosis (*rho* = −0.25, *P* = 0.001, Supplementary Figure S3), this association was even stronger when restricting to ER positive patients (*rho* = −0.37, *P* < 0.001, Supplementary Figure S3). In a linear regression analysis adjusted for potential cofounders including pathological stage, molecular subtypes by IHC and histological subtypes we found that age at first birth was independently and negatively associated with age at diagnosis (*P* = 0.0004).

**Table 1.** Clinicopathological features of nulliparous and parous patients.

|  |  | **Nulliparous** | **Parous** | **P** |  | **Early parous** | **Late parous** | **P** |
| --- | --- | --- | --- | --- | --- | --- | --- | --- |
| **N** |  | **49** | **264** |  |  | **82** | **71** |  |
| **Age** |  | 54 (30-81) | 55 (28-81) | 0.796<sup>a</sup> |  | 59 (34-81) | 49 (28-81) | 2.6x10<sup>-5</sup> <sup>a</sup> |
| **Menopausal status** | **Pre** | 13 (33.3%) | 60 (30.3%) |  |  | 14 (20.9%) | 29 (45.3%) |  |
|  | **Post** | 26 (66.7%) | 138 (69.7%) | 0.85 |  | 53 (79.1%) | 35 (54.7%) | 0.0053 |
| **Stage** | **I** | 7 (14.9%) | 41 (15.9%) |  |  | 6 (7.6%) | 17 (24.3%) |  |
|  | **II** | 20 (42.6%) | 81 (31.4%) |  |  | 33 (41.8%) | 27 (38.6%) |  |
|  | **III** | 11 (23.4%) | 30 (11.6%) |  |  | 12 (15.2%) | 10 (14.3%) |  |
|  | **IV** | 0 (0%) | 1 (0.4%) |  |  | 1 (1.3%) | 0 (0%) |  |
|  | **Na** | 9 (19.1%) | 105 (40.7%) | 0.019 |  | 27 (34.2%) | 16 (22.9%) | 0.039 |
| **pT** | **Tx** | 9 (18.4%) | 105 (39.8%) |  |  | 27 (32.9%) | 16 (22.5%) |  |
|  | **<2cm** | 11 (22.4%) | 61 (23.1%) |  |  | 13 (15.9%) | 26 (36.6%) |  |
|  | **>2cm** | 29 (59.2%) | 98 (37.1%) | 0.0062 |  | 42 (51.2%) | 29 (40.8%) | 0.012 |
| **pN** | **Nx** | 11 (22.4%) | 112 (42.4%) |  |  | 30 (36.6%) | 17 (23.9%) |  |
|  | **N0** | 18 (36.7%) | 79 (29.9%) |  |  | 25 (30.5%) | 23 (32.4%) |  |
|  | **N1+** | 20 (40.8%) | 73 (27.7%) | 0.027 |  | 27 (32.9%) | 31 (43.7%) | 0.2 |
| **Grade** | **I** | 6 (14%) | 29 (12.7%) |  |  | 8 (9.8%) | 5 (7%) |  |
|  | **II** | 17 (39.5%) | 86 (37.7%) |  |  | 25 (30.5%) | 35 (49.3%) |  |
|  | **III** | 20 (46.5%) | 113 (49.6%) | 0.93 |  | 49 (59.8%) | 31 (43.7%) | 0.059 |
| **Subtype by IHC** | **Lum A-like** | 22 (44.9%) | 106 (40.3%) |  |  | 29 (35.4%) | 39 (54.9%) |  |
|  | **Lum B-like** | 19 (38.8%) | 54 (20.5%) |  |  | 20 (24.4%) | 17 (23.9%) |  |
|  | **HER2+/HR+** | 6 (12.2%) | 29 (11%) |  |  | 0 (0%) | 0 (0%) |  |
|  | **HER2+/HR-** | 0 (0%) | 13 (4.9%) |  |  | 1 (1.2%) | 1 (1.4%) |  |
|  | **TNBC** | 2 (4.1%) | 61 (23.2%) | 0.0013 |  | 32 (39%) | 14 (19.7%) | 0.026 |
| **Histology** | **Ductal** | 36 (76.6%) | 203 (81.2%) |  |  | 71 (88.8%) | 49 (71%) |  |
|  | **Lobular** | 5 (10.6%) | 23 (9.2%) |  |  | 2 (2.5%) | 10 (14.5%) |  |
|  | **Other** | 6 (12.8%) | 24 (9.6%) | 0.67 |  | 7 (8.8%) | 10 (14.5%) | 0.01 |

pT, pathological tumor size; pN, pathological nodal status; HR, hormone receptor; P, p-value derived from $\chi^2$ test or the Fisher exact test when appropriate (<sup>a</sup>except continuous variable derived from Mann–Whitney U test)

In the following sections, we investigate the imprint of parity and age at first birth on breast cancer biology by using a multivariate analysis adjusted for potential cofounders, namely age at diagnosis, pathological stage, molecular subtypes by IHC and histological subtypes.

### The influence of parity and age at first birth on breast cancer somatic mutational load

We first sought to investigate the imprint of parity and age at first birth on somatic mutational load. There was no significant difference in the total number of substitutions (SNVs) according to parity nor age at first birth (*P*_*adj*_ = 0.097, *P*_*adj*_ = 0.075, respectively, Figure 1a). There was no significant difference in the total number of insertions or deletions (Indels) according to parity (*P*_*adj*_ = 0.464, Figure 1a). In contrary, compared to tumors from late parous patients, tumors from early parous patients were significantly associated with a higher Indels load (*P*_*adj*_ = 0.002, *FDR* = 0.007, Figure 1a). There was no significant difference between the total number of rearrangements according to parity nor age at first birth (Supplementary Table S1).

**Figure 1.**
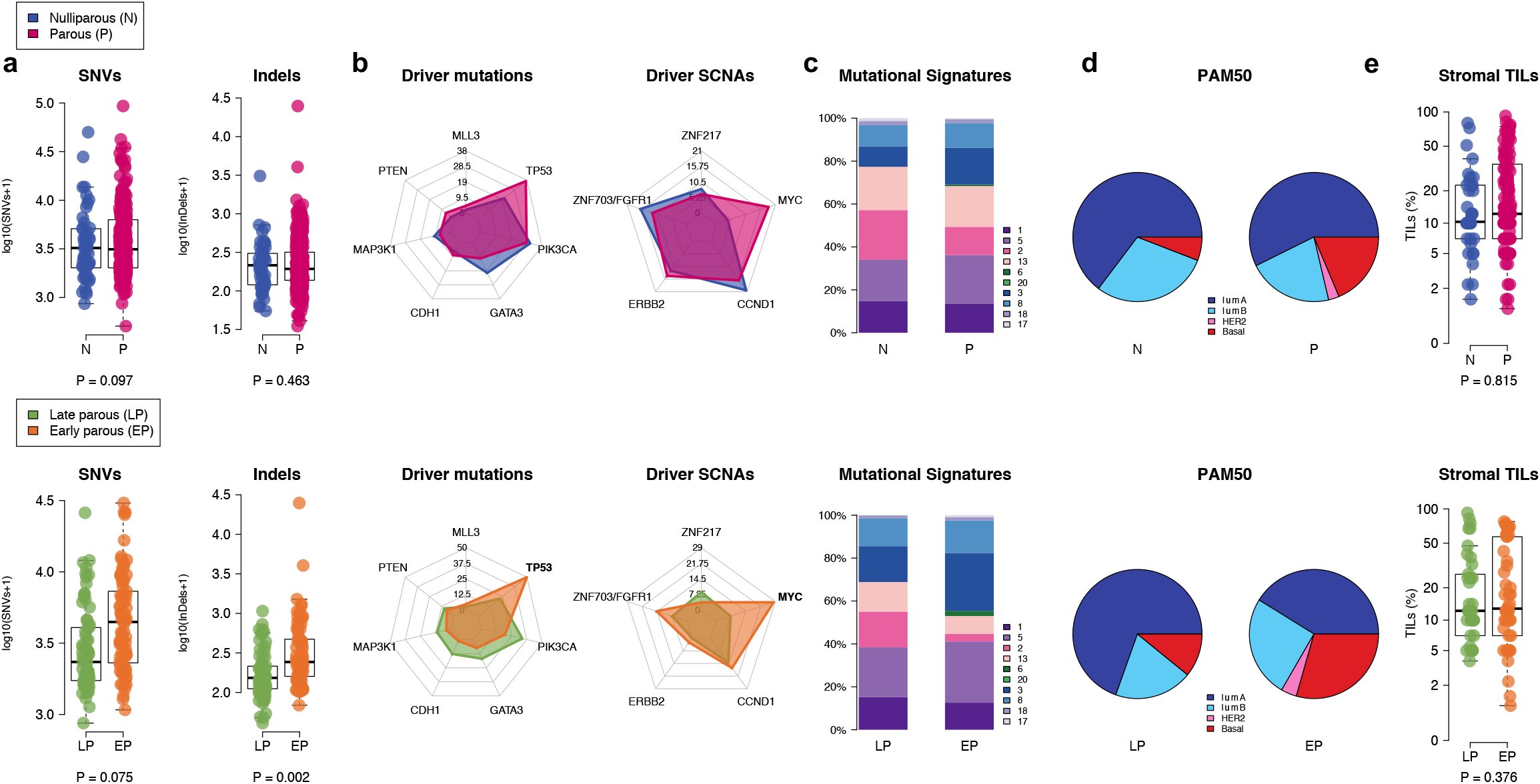
Imprint of pregnancy and age at first birth on breast cancer biology. (a) Comparison of SNVs and Indels between nulliparous and parous (upper) and between early and late parous (bottom). P-values are derived from multivariate linear regression analysis adjusted for potential confounders. (b) Radar plots showing the frequency of driver mutations and driver SNCAs between nulliparous and parous (upper) and between early and late parous (bottom) patients. Significant genes independently associated with parity or age at first birth are highlighted in bold. (c) Proportion of breast cancer substitution signatures in nulliparous and parous (upper) and in early and late parous (bottom) patients. (d) Proportion of PAM50 breast cancer subtypes in nulliparous and parous (upper) and in early and late parous (bottom) patients. (e) Comparison of TILs levels (%) between nulliparous and parous (upper) and between early and late parous (bottom) patients. P-values are derived from multivariate linear regression analysis adjusted for potential confounders.

We next interrogated the influence of parity and age at first birth on the frequency of mutations in breast cancer driver genes. Among the driver mutated genes, seven had at least one non-silent mutation with a frequency of > 5% across the whole cohort. As expected, *PIK3CA* and *TP53* were the most frequently mutated genes (Figure 1b). None of the driver mutated genes were associated with parity in the multivariate analysis (Supplementary Table S1). However, in the parous group, early age at first birth was independently associated with higher frequency of *TP53* mutations (50% vs. 22.5%; *P*_*adj*_ = 0.010; *FDR* = 0.046, Figure 1b) and lower frequency of *CDH1* mutations (1.2% vs. 12.7%; *P*_*adj*_ = 0.013; *FDR* = 0.046, Figure 1b). Considering the distribution of *TP53* mutations type, early parous patients had a significantly higher frequency of truncating mutations as compared to late parous patients (25.6% vs. 7%; *P*_*adj*_ = 0.014, Supplementary Figure S4). Altogether, our results show that age at first birth is associated with biological differences in the mutational landscape of subsequent breast tumours with early parity associated with higher Indels burden and higher frequency of deleterious mutations in *TP53* gene.

### The influence of parity and age at first birth on somatic copy number alterations

Somatic copy number alterations (SCNAs) play a major role in breast cancer biology (14,15). We identified 5 driver genes with a frequency of SCNAs > 5% across all patients. *MYC* tended to be more frequently amplified in parous than in nulliparous patients (18.6% vs. 4.1 %; *P*_*adj*_ = 0.052; *FDR* = 0.26, Figure 1 b). In the parous group, *MYC* amplification was significantly more frequent in the early parous group than in the late parous group (28% vs. 7%; *P*_*adj*_ = 0.008; FDR = 0.040, Figure 1b). When evaluating the co-occurrence of SCNAs and somatic mutations, we found that co-occurrence of *MYC* amplification and *TP53* mutations was independently associated with age at first birth, with early parous patients having a higher frequency of simultaneous alterations of *MYC* and *TP53* genes (18.3% vs. 4.2%; *P*_*adj*_ = 0.087, Figure 2a and Supplementary Figure S4). Taken together, these results suggest that age at first birth may also shape the somatic copy number alterations profiles of subsequent breast cancer.

**Figure 2.**
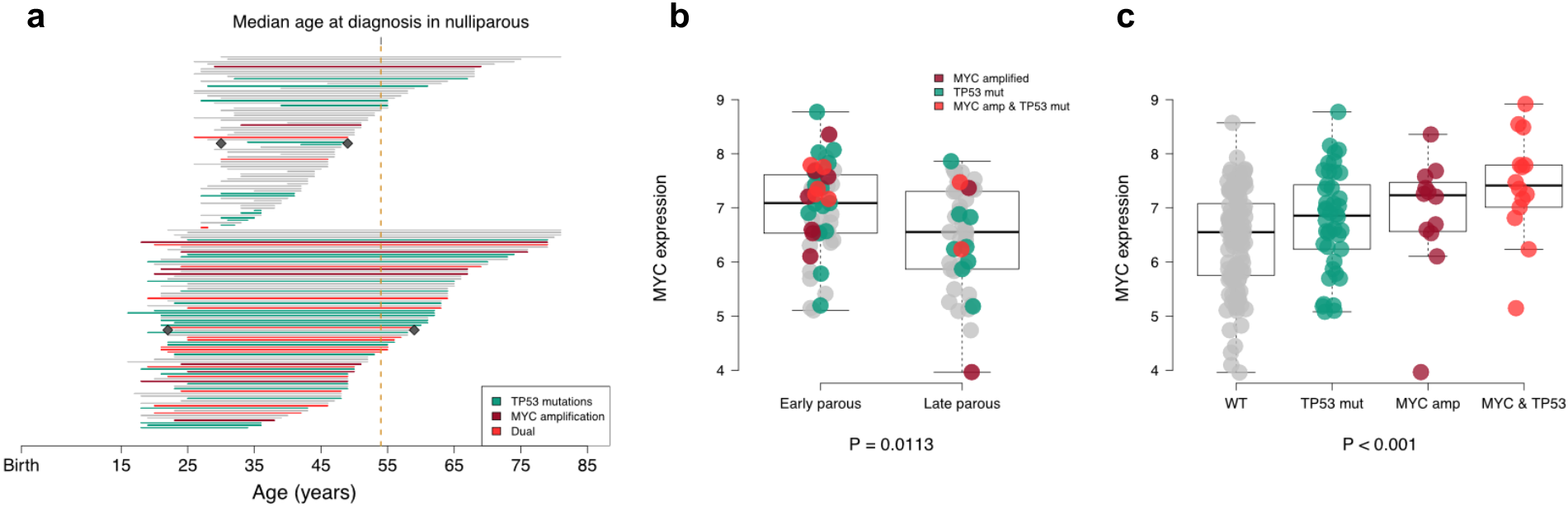
Co-occurrence of *MYC* amplification and *TP53* mutations is associated with age at first birth. (a) Timeline of 153 patients with available data on age at first birth. Each line represents an individual patient from age at first birth (start of the line) to age at breast cancer diagnosis (end of the line). Late parous patients (upper) and early parous patients (bottom) are grouped according to median age at first birth. Grey diamond represents the median age at first birth and at diagnosis in the two groups. Lines are colored according to *TP53* mutations (green) *MYC* amplification (dark red) and the co-occurrence of both (red). (b) Comparison of *MYC* expression in early and late parous patients. Each dot represents an individual patient and is colored according to *TP53* mutations (green) *MYC* amplification (dark red) and the co-occurrence of both (red). P-value is derived from multivariate linear regression analysis adjusted for potential confounders. (c) *MYC* expression according to *TP53* mutations, *MYC* amplification or the co-occurrence of both. P-value is derived from Kruskal-Wallis test.

### The influence of parity and age at first birth on mutational signatures

To have a better understanding of the mutational processes that have occurred during the course of cancer according to reproductive history, we examined the contribution of mutational substitution signatures known to occur in breast cancer (13). The distribution of mutational signatures was similar between parous and nulliparous patients. However, in the parous group, signatures 2 was more prevalent in late parous patients (*P*_*adj*_ = 0.001; *FDR* = 0.011, Figure 1c). Of interest, two early parous patients with exceptionally high number of Indels were associated with mutational processes 6 and 26, attributable to mismatch repair deficiency (16) (Supplementary Figure S4). Similarly to our previous findings, these results highlight that age at first birth may be associated with specific mutational processes in subsequent breast cancer.

### The influence of parity and age at first birth on BRCAness

We investigated the BRCAness status of tumors according to reproductive history. Since germline BRCA1/2 mutation status was not available for all samples, we used HRDetect to identify *BRCA1/BRCA2-deficient* samples (17). We did not find any significant differences in the proportion of BRCA1/BRCA2-deficient patients between nulliparous and parous groups (12.24% vs. 18.94%, respectively, *P*_*adj*_ = 0.473), nor between early and late parous group (30.49% vs. 19.71%, respectively, *P*_*adj*_ = 0.386). Thus, reproductive history and age at first birth do not seem to affect homologous recombination DNA repair capacity in subsequent breast cancer.

### Integrative analysis of the genomic alterations and the transcriptomic profiles associated with parity and age at first birth

RNA sequencing data was available for a subset of 182 patients, of which 34 were nulliparous (Supplementary Figure S2 and Table S2). We first determined the intrinsic molecular subtypes distribution of breast cancer using the PAM50 classifier (18). We did not find a significant difference in the distribution of the PAM50 subtypes between nulliparous and parous (Figure 1d). In contrast, early parous patients had a higher proportion of basal-like subtype tumors (29.4% vs. 8.9%; *P* = 0.009, Figure 1d). In order to identify *de novo* gene expression profiles that might be associated with parity and age at first birth, we performed a multivariate differential expression analysis using DEseq2 (19) controlling for age at diagnosis, pathological stage, molecular subtypes by IHC and histological subtypes. A total of 62 genes were differentially expressed between nulliparous and parous (Supplementary Table S3). Among these genes, three were associated with mammary development. *OXTR* and *ATP2B2* were down-regulated whereas *NRG3* was up-regulated in nulliparous. Pathway analysis using the generally applicable gene-set enrichment (GAGE) analysis (20) revealed an enrichment of genes related to extracellular matrix (ECM) receptor interaction (Supplementary Table S4). When comparing early and late parous patients, 466 genes were differentially expressed, among which 305 were up-regulated in early parous (Supplementary Table S5). However, pathway analysis did not reveal any significant enrichment of relevant biological processes (Supplementary Table S6).

Due to the higher frequency of *MYC* amplification in early parous patients, we determined if this would also impact *MYC* at the mRNA expression levels. Early parous patients were independently associated with an up-regulation of *MYC* expression (*P*_*adj*_ = 0.0113, Figure 2b). We also evaluated the expression of *MYC* according to *TP53* mutations and *MYC* amplification and found that *MYC* expression was the highest in tumors harboring concurrent *TP53* mutation and *MYC* amplification (Figure 2c).

Signature 2 and 13 have been attributed to activity of the AID/APOBEC family of cytidine deaminases converting cytosine to uracil (16). Thus, we inspected if the higher prevalence of signature 2 observed in late parous patients was associated with higher expression of AID/APOBEC expression. As opposed to signature 13, which was positively correlated with all APOBEC3s family members and AID expression, signature 2 was not significantly associated with the expression of any of these deaminases (Supplementary Figure S5).

### The influence of parity and age at first birth on immune infiltration

Previous reports have hypothesized that the pregnancy-induced tumor protection could be attributable to an improved anti-tumor immunity (21–25). Therefore, we assessed whether reproductive history could be associated with tumor infiltrating lymphocyte (TILs) levels that is considered as a surrogate of tumor immunogenicity. We did not find any significant difference in the proportion of stromal TILs according to parity or according to the age at first birth (*P*_*adj*_ = 0.644; *P*_*adj*_ = 0.376, respectively, Figure 2e). Similarly, no differences were observed when comparing intratumoral TILs according to parity or age at first birth (*P*_*adj*_ = 0.242; *P*_*adj*_ = 0.993). Thus, reproductive history does not seem to influence breast tumour immunogenicity.

### Pregnancy associated breast cancer are associated with increased TILs infiltration

Pregnancy associated breast cancer (PABC) can be defined as cases diagnosed up to 10 years postpartum (26). In this cohort, we identified 17 PABC patients and compared them with nulliparous patients. Compared to nulliparous, PABC patients had a younger age at diagnosis (median, 38 years; range, 28-48 years vs. median, 54 years; range, 30-81 years; *P* = 5.79 × 10^−6^, Supplementary Table S7) and higher frequency of TNBC (29.4% vs. 4.1%; *P* = 0. 021, Supplementary Table S7). We did not find any significant differences in the pattern of somatic mutations, somatic copy number alterations (SCNAs) nor in the distribution of mutational signatures (Supplementary Table S1). Nine PABC had available gene expression and TILs scoring. At the transcriptomic level, we found that PABC patients were associated with enrichment of biological processes related to immune function (Figure 3a). Moreover, PABC patients had an increased lymphocytic infiltration both for stromal and intratumoral TILs levels (*P*_*adj*_ = 0.0495 and *P*_*adj*_ = 0.00003, respectively; Figure 3bc). Taken together these results indicate that cancer occurring in the postpartum mammary gland is associated with an increased immune function.

**Figure 3.**
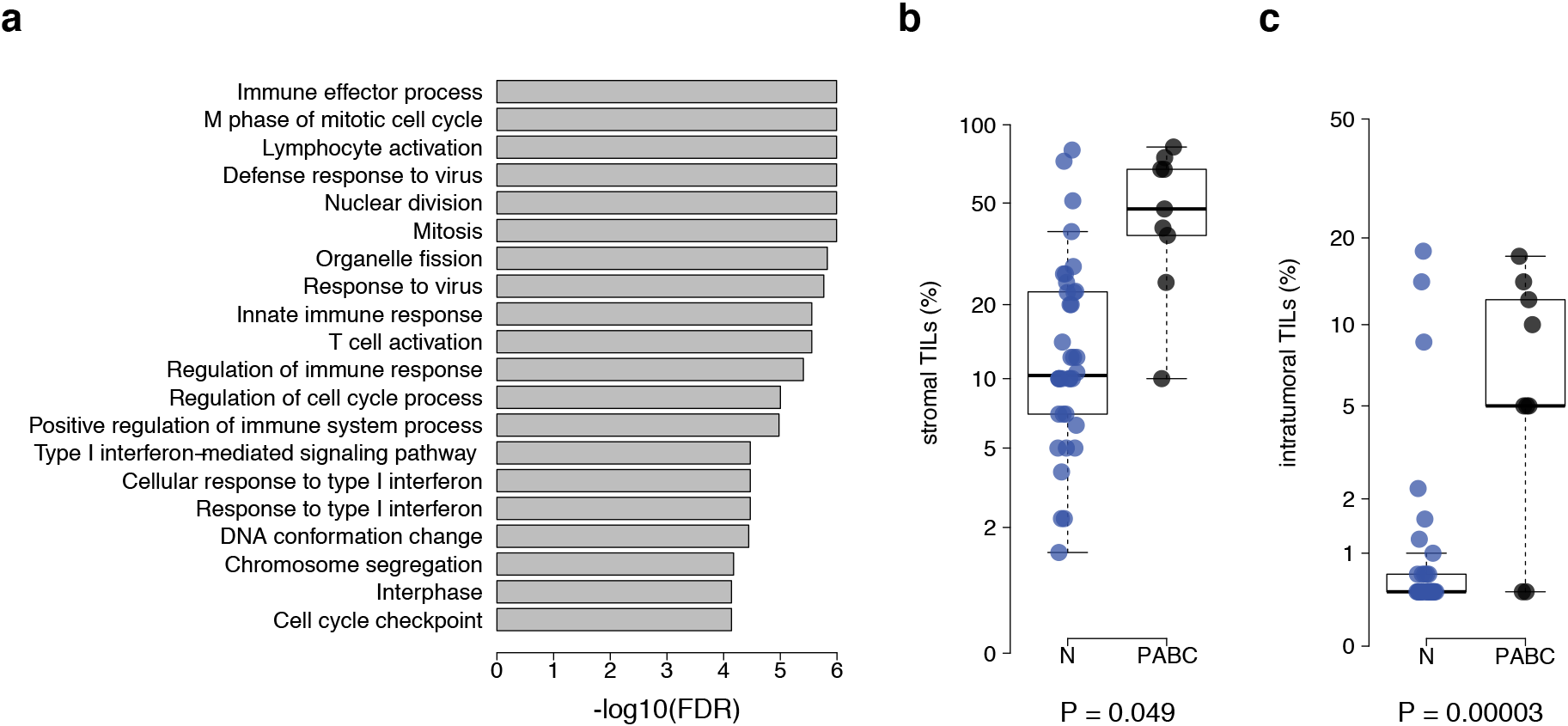
PABC patients are associated with higher TILs levels. (a) Results form the GAGE analysis showing the top 20 most significant biological processes enriched in PABC patients. (b) Comparison of stromal and (c) intratumoral (right) TILs levels (%) between nulliparous (N) and PABC. P-values are derived from multivariate linear regression analysis adjusted for potential confounders.

## Discussion

To our knowledge, this is the first study that explores the impact of reproductive history on the genomic landscape and the immune composition of subsequent breast cancer. While previous studies documented the risk of developing breast cancer according to reproductive history (1–5), this analysis provides further insights on the differences at the pathologic, genomic, transcriptomic and immunogenic levels according to prior parity and age at first birth. Independently of clinicopathological features, our findings indicate that age at first birth impacts the genomic makeup of subsequent breast cancer. Early parous patients developed tumors characterized by a higher number of Indels, a lower frequency of *CDH1* mutations, a higher frequency of *TP53* mutations and *MYC* amplification and a lower prevalence of mutational signature 2, while PACB patients exhibited higher TILs infiltration.

The higher proportion of TNBC in parous and particularly in early parous patients could be attributed to a differential effect of pregnancy-induced tumor protection according to breast cancer subtypes. We and others have documented that the pregnancy-induced tumor protection is different according to breast cancer subtypes with parity and young age at first birth being associated with a marked reduction in the risk of developing luminal subtype (7–10). For the first time to the best of our knowledge, we have documented that age at first birth is negatively associated with age at diagnosis irrespective of classical clinicopathological features. This observation, that needs to be validated in larger cohorts, is in line with the reported protective effect of an early pregnancy on breast cancer risk.

Our study reveals that age at first birth has a bigger imprint on genomic alterations of breast cancer than parity status alone. However, the apparent lack of impact of parity could be also related to the relatively low number of nulliparous patients.

At the gene level, early parous patients had a higher frequency of *TP53* mutation, *MYC* amplification and a lower frequency of *CDH1*. Interestingly, co-occurrence of *TP53* mutations and *MYC* amplification was independently associated with age at first birth, while the proportion of truncating *TP53* mutations was higher in early parous patients. We observed that tumors harboring concurrent *MYC* amplification and *TP53* mutation had the highest *MYC* expression. This observation is in line with a recent investigation of the *MYC* oncogene in pan-cancer data (27). Previous reports have suggested that *TP53* mutations are a common mechanism that disturb the apoptotic pathway in MYC-driven tumors (28). It has been hypothesized that overexpression of *MYC* induces *TP53*-dependent apoptosis, and, as a consequence, MYC-driven tumors often require dysregulation of the apoptotic pathway to promote proliferation (29). *TP53* has long been recognized as a potential mediator of pregnancy-induced resistance to mammary carcinogenesis. It has been shown that p53 and its downstream transcriptional target p21, are increased in parous and estrogen/progesterone-treated mammary epithelium in response to carcinogen (30). In the absence of p53, the protection given by parity or exogenous hormones is lost (31–33). We hypothesized that the higher frequency of *TP53* mutation observed in breast cancer from early parous woman could be explained by the fact that an early pregnancy might protect less effectively against *TP53* mutated breast cancer. In breast cancer, *TP53* mutations are highly linked to molecular subtype with a frequency of 80% in basal-like compared to 26% in luminal tumors (15). The differential effect of parity-induced protection according to *TP53* mutational status might also explain the differential effect of parity-induced protection according to tumor subtypes.

*CDH1* mutations has been associated with invasive lobular breast cancer subtype (34). As the multivariate analysis was adjusted for histological subtypes, the lower frequency of *CDH1* mutations observed in early parous patients cannot be explained by differences in histological subtypes. The lower level of mutational signature 2 seen in early parous is not explained by difference in the expression of AID/APOBEC family but could be related to other factors that remained to be determined.

At the mRNA level, we found that age at first birth had a stronger impact on the transcriptome than parity status alone, indicating again that age at first birth might be the most critical factor. In the normal tissue of parous women, the gene encoding oxytocin receptor (*OXTR*) is physiologically up-regulated during lactation and has been shown to remain overexpressed later in life (12,35). Noteworthy, the expression of *OXTR* was higher in parous compared to nulliparous patients. However, due to the lack of functional studies, it is not clear whether this gene is involved in the tumorigenesis of breast cancer or simply related to physiological changes induced by pregnancy. The enrichment of genes related to ECM receptor interaction in parous patients might be related to involution, a profound physiological change in the mammary gland after pregnancy. Right after breastfeeding the fully differentiated gland regresses to its pre-pregnant state by an innate tissue-remodelling mechanism. Evidence indicates that involution is mediated in part by ECM-degrading proteinases, leading to basement-membrane degradation and subsequent apoptosis of the unwanted secretory epithelial cells (36,37). The exact role of the enrichment of genes related to ECM in parous patients on human breast cancer biology has still to be determined, but involution, that shares similarities with inflammation and wound healing programs, has been shown to promote breast cancer progression and metastasis in several animal models (36,38).

Finally, previous reports have hypothesized that the pregnancy-induced tumor protection could be attributable to an improved anti-tumor immunity (21–25). Our analysis reveals no differences in TILs infiltration levels according to parity or age at first birth. The existence of a more complex immune component related to reproductive history cannot be excluded, but it is not supported by our analysis. Previous reports have documented that the immune milieu of the postpartum mammary gland caused by involution could contribute to tumor promotion (39–41). We observed an increase of TILs levels in PABC patients but the composition of the immune infiltrate has still be determined to validate this hypothesis.

A potential limitation of our study is the lack of data on other reproductive factors (e.g. breastfeeding, age at menarche and time since last birth) that could also potentially imprint the genomic alterations of breast cancer. Indeed, breastfeeding and age at menarche have been also linked to breast cancer risk but since they are often self-reported, they are more difficult to assess reliably (7). Another limitation is the absence of HER2+ subtype in the transcriptomic analysis.

## Conclusions

In conclusion, our findings highlight an unprecedented link between reproductive factors and the genomic landscape of subsequent breast cancer. Specifically, our analysis suggests that age at first birth, a known breast cancer risk factor, adds a layer of biological complexity to subsequent breast tumors. Our results, that need to be validated in other studies, support that patients’ reproductive history should be routinely collected in future large scale genomic studies addressing the biology of female cancers. With the rapid development of precision oncology, this work also advocates that reproductive history should not be underestimated in future clinical studies of breast cancer.

## Methods

### Data acquisition

All analyses were performed on a publicly available dataset comprising 560 breast cancer patients referred to as BRCA560 (13). Clinical data, sequencing coverage and mutational load were obtained from Supplementary Tables 1-3 in that reference. Coding driver mutation events and the contribution of mutational signatures were obtained from Supplementary Tables 14 and 21 in that reference. Raw count data from RNA sequencing were obtained from the authors. Results from HRDetect classifier were obtained from Supplementary Table 4 in reference (17).

### Patients selection

Eligible patients from BRCA560 were those with samples collected from primary tumor only (patients with local recurrence or metastasis samples were excluded, N = 8) who had available information on parity. There were only two available HER2+ patients (both parous) in the transcriptomic analysis so we preferred to exclude them from this analysis. For each patient, we determined the breast cancer intrinsic subtype using PAM50 (18). PAM50 classes were determined from the BRCA560 RNA-seq gene expression data using the genefu R/Bioconductor package (42). Nulliparous patients were defined as women with breast cancer who have never given birth. Parous patients were defined as women with breast cancer who had at least one full term pregnancy. Early parous patients were defined as ≤ 25 years of age at first full term pregnancy, while late parous patients were defined as > 25 years of age at first full term pregnancy. Since the BRCA560 dataset is publicly available, ethics committee approval was not needed. In addition, neither patient informed consent nor permission to use these data were required to perform this analysis.

### Statistical analysis

Except for age at diagnosis that was considered as a continuous variable and therefore compared using the non-parametric Mann-Whitney U test, differences in other clinicopathological characteristics of breast cancer between groups were analysed using the χ2 test or the Fisher exact test when appropriate. All statistical tests comparing groups were done using the non-parametric Mann-Whitney U test and the Fisher exact test for continuous and categorical variables, respectively. For the multivariate analysis, we used a linear and logistic regression to assess the independent association of continuous (log transformed) and categorical variables respectively with – parity (nulliparous vs parous) or – age at first birth (≤ 25 years vs. > 25 years) controlling for: age at diagnosis, pathological stage, molecular subtypes by IHC, histological subtypes. For WGS results, we also corrected for log-transformed sequence coverage of tumor and normal samples (continuous). All interaction and multivariate tests (*P*_*adj*_) were done using analysis of variance to compare the models with and without the extra term. Because continuous variables contain zeros, the logarithmic transformation was applied as follows: log_10_(x + 1). The Kruskal-Wallis test was used to test if *MYC* expression originates from the same population according to genomic status of *MYC/TP53* alterations. All correlations were measured using the non-parametric Spearman’s *rho* coefficient. All reported P-values were two-tailed. Multiple testing correction was done using the false discovery rate method (FDR) (43) and differences were considered significant when the FDR was < 0.05. All analyses were done in R software version 3.3.3 (available at www.r-project.org) and Bioconductor version 3.6. Differential expression analysis was performed with DESeq2 v.1.14.1 R/Bioconductor package (19) on raw count data. Significantly differentially expressed genes were selected with a false discovery rate (FDR) of < 0.1. We used gage v.2.24.0 R/Bioconductor package (20) to identify significantly enriched Kyoto Encyclopaedia of Genes and Genomes (KEGG) pathways (44) with the log2FoldChange from DEseq2 results as input data.

### TILs evaluation

The percentage of TILs was independently evaluated by two pathologists (R.S. and H.H.) on hematoxylin and eosin slides using the International TILs Working Group 2014 methodology as described before (45). There were 242 original samples with evaluable TILs from 239 patients. For the three patients with two samples the arithmetic averages were obtained. We obtain a final set of 231 patients with primary tumor only (patients with local recurrence or metastasis samples only were excluded, N = 8). TILs information for patients for which evaluation from only one pathologist was available were discarded (N=3).

## References

1. MacMahon B, Cole P, Lin TM, Lowe CR, Mirra AP, Ravnihar B, et al. Age at first birth and breast cancer risk. Bull World Health Organ. 1970;43(2):209–21.

2. Papatestas AE, Mulvihill M, Josi C, Ioannovich J, Lesnick G, Aufses AH. Parity and prognosis in breast cancer. Cancer. 1980 Jan 1;45(1):191–4.

3. Trichopoulos D, Hsieh CC, MacMahon B, Lin TM, Lowe CR, Mirra a P, et al. Age at any birth and breast cancer risk. Int J Cancer. 1983;31(6):701–4.

4. Kroman N, Wohlfahrt J, Andersen KW, Mouridsen HT, Westergaard T, Melbye M. Time since childbirth and prognosis in primary breast cancer: population based study. BMJ. 1997 Oct 4;315(7112):851–5.

5. Rosenberg L, Thalib L, Adami H-O, Hall P. Childbirth and breast cancer prognosis. Int J Cancer. 2004 Sep 20;111 (5):772–6.

6. Albrektsen G, Heuch I, Hansen S, Kvåle G. Breast cancer risk by age at birth, time since birth and time intervals between births: exploring interaction effects. Br J Cancer. 2005;92(1):167–75.

7. Lambertini M, Santoro L, Del Mastro L, Nguyen B, Livraghi L, Ugolini D, et al. Reproductive behaviors and risk of developing breast cancer according to tumor subtype: A systematic review and meta-analysis of epidemiological studies. Cancer Treat Rev. Elsevier Ltd; 2016;49:65–76.

8. Yang XR, Chang-Claude J, Goode EL, Couch FJ, Nevanlinna H, Milne RL, et al. Associations of breast cancer risk factors with tumor subtypes: A pooled analysis from the breast cancer association consortium studies. J Natl Cancer Inst. 2011;103(3):250–63.

9. Ellingjord-Dale M, Vos L, Tretli S, Hofvind S, dos-Santos-Silva I, Ursin G. Parity, hormones and breast cancer subtypes - results from a large nested case-control study in a national screening program. Breast Cancer Res. Breast Cancer Research; 2017;19(1):10.

10. Ritte R, Tikk K, Lukanova A, Tjønneland A, Olsen A, Overvad K, et al. Reproductive factors and risk of hormone receptor positive and negative breast cancer: A cohort study. BMC Cancer. 2013;13:1–12.

11. Meier-Abt F, Bentires-Alj M. How pregnancy at early age protects against breast cancer. Trends Mol Med. Elsevier Ltd; 2014;20(3):143–53.

12. Russo J, Santucci-Pereira J, De Cicco RL, Sheriff F, Russo PA, Peri S, et al. Pregnancy-induced chromatin remodeling in the breast of postmenopausal women. Int J Cancer. 2012;131(5):1059–70.

13. Nik-Zainal S, Davies H, Staaf J, Ramakrishna M, Glodzik D, Zou X, et al. Landscape of somatic mutations in 560 breast cancer whole-genome sequences. Nature. Nature Publishing Group; 2016;534(7605):1–20.

14. Curtis C, Shah SP, Chin S-F, Turashvili G, Rueda OM, Dunning MJ, et al. The genomic and transcriptomic architecture of 2,000 breast tumours reveals novel subgroups. Nature. 2012;486(7403):346–52.

15. TCGA. Comprehensive molecular portraits of human breast tumours. Nature. 2012;487(7407):61–70.

16. Alexandrov LB, Nik-Zainal S, Wedge DC, Aparicio S a JR, Behjati S, Biankin A V, et al. Signatures of mutational processes in human cancer. Nature. 2013;500:415–21.

17. Davies H, Glodzik D, Morganella S, Yates LR, Staaf J, Zou X, et al. HRDetect is a predictor of BRCA1 and BRCA2 deficiency based on mutational signatures. Nat Med (in Press. 2017;(November 2016).

18. Parker JS, Mullins M, Cheang MCU, Leung S, Voduc D, Vickery T, et al. Supervised Risk Predictor of Breast Cancer Based on Intrinsic Subtypes. J Clin Oncol. 2009 Mar 10;27(8):1160–7.

19. Love MI, Huber W, Anders S. Moderated estimation of fold change and dispersion for RNA-seq data with DESeq2. Genome Biol. 2014 Dec 5;15(12):550.

20. Luo W, Friedman MS, Shedden K, Hankenson KD, Woolf PJ. GAGE: generally applicable gene set enrichment for pathway analysis. BMC Bioinformatics. BioMed Central; 2009 May 27;10(1):161.

21. Erlebacher A. Mechanisms of T cell tolerance towards the allogeneic fetus. Nat Rev Immunol. Nature Publishing Group; 2013 Jan 14;13(1):23–33.

22. Agrawal B, Reddish MA, Krantz MJ, Longenecker BM. Does pregnancy immunize against breast cancer? Cancer Res. 1995 Jun 1;55(11):2257–61.

23. Finn OJ, Jerome KR, Henderson RA, Pecher G, Domenech N, Magarian-Blander J, et al. MUC-1 epithelial tumor mucin-based immunity and cancer vaccines. Immunol Rev. 1995 Jun;145:61–89.

24. Jungbluth AA, Silva WA, Iversen K, Frosina D, Zaidi B, Coplan K, et al. Expression of cancer-testis (CT) antigens in placenta. Cancer Immun. 2007 Aug 24;7:15.

25. Arklie J, Taylor-Papadimitrious J, Bodmer W, Egan M, Millis R. Differentiation antigens expressed by epithelial cells in the lactating breast are also detectable in breast cancers. Int J cancer. 1981 Jul 15;28(1):23–9.

26. Lyons TR, Schedin PJ, Borges VF. Pregnancy and breast cancer: When they collide. J Mammary Gland Biol Neoplasia. 2009;14(2):87–98.

27. Leiserson MDM, Vandin F, Wu HT, Raphael BJ. Reply: Co-occurrence of MYC amplification and TP53 mutations in human cancer. Nat Genet. 2016;48(2):106–8.

28. Wolf E, Lin CY, Eilers M, Levens DL. Taming of the beast: Shaping Myc-dependent amplification. Trends Cell Biol. Elsevier Ltd; 2015;25(4):241–8.

29. Hermeking H, Eick D. Mediation of c-Myc-induced apoptosis by p53. Science (80-). 1994;265(5181):2091–3.

30. Sivaraman L, Conneely OM, Medina D, O’Malley BW. P53 Is a Potential Mediator of Pregnancy and Hormone-Induced Resistance To Mammary Carcinogenesis. Pnas. 2001;98(22):12379–84.

31. Dunphy KA, Blackburn AC, Yan H, O’Connell LR, Jerry DJ. Estrogen and progesterone induce persistent increases in p53-dependent apoptosis and suppress mammary tumors in BALB/c-Trp53+/−mice. Breast Cancer Res. BioMed Central; 2008 Jun 12;10(3):R43.

32. Medina D, Kittrell FS. p53 function is required for hormone-mediated protection of mouse mammary tumorigenesis. Cancer Res. 2003 Oct 1; 63(19):6140–3.

33. Jerry DJ, Kittrell FS, Kuperwasser C, Laucirica R, Dickinson ES, Bonilla PJ, et al. A mammary-specific model demonstrates the role of the p53 tumor suppressor gene in tumor development. Oncogene. 2000 Feb 7;19(8):1052–8.

34. Desmedt C, Zoppoli G, Gundem G, Pruneri G, Larsimont D, Fornili M, et al. Genomic Characterization of Primary Invasive Lobular Breast Cancer. J Clin Oncol. 2016;34(16):1872–80.

35. Peri S, de Cicco RL, Santucci-Pereira J, Slifker M, Ross EA, Russo IH, et al. Defining the genomic signature of the parous breast. BMC Med Genomics. 2012;5:46.

36. McDaniel SM, Rumer KK, Biroc SL, Metz RP, Singh M, Porter W, et al. Remodeling of the mammary microenvironment after lactation promotes breast tumor cell metastasis. Am J Pathol. 2006 Feb;168(2):608–20.

37. Schedin P. Pregnancy-associated breast cancer and metastasis. Nat Rev Cancer. 2006;6(4):281–91.

38. Lyons TR, O’Brien J, Borges VF, Conklin MW, Keely PJ, Eliceiri KW, et al. Postpartum mammary gland involution drives progression of ductal carcinoma in situ through collagen and COX-2. Nat Med. Nature Research; 2011 Aug 7;17(9):1109–15.

39. Martinson HA, Jindal S, Durand-Rougely C, Borges VF, Schedin P. Wound healing-like immune program facilitates postpartum mammary gland involution and tumor progression. Int J Cancer. 2015 Apr 15;136(8):1803–13.

40. Fornetti J, Martinson HA, Betts CB, Lyons TR, Jindal S, Guo Q, et al. Mammary Gland Involution as an Immunotherapeutic Target for Postpartum Breast Cancer. J Mammary Gland Biol Neoplasia. 2014 Jul 22;19(2):213–28.

41. Harvell DME, Kim J, O’Brien J, Tan A-C, Borges VF, Schedin P, et al. Genomic signatures of pregnancy-associated breast cancer epithelia and stroma and their regulation by estrogens and progesterone. Horm Cancer. NIH Public Access; 2013 Jun;4(3):140–53.

42. Gendoo DMA, Ratanasirigulchai N, Schröder MS, Paré L, Parker JS, Prat A, et al. Genefu: an R/Bioconductor package for computation of gene expression-based signatures in breast cancer. Bioinformatics. 2016 Apr 1;32(7):1097–9.

43. Benjamini Y, Hochberg Y. Controlling the false discovery rate: a practical and powerful approach to multiple testing. Vol. 57, Journal of the Royal Statistical Society B. 1995. p. 289–300.

44. Kanehisa M, Furumichi M, Tanabe M, Sato Y, Morishima K. KEGG: new perspectives on genomes, pathways, diseases and drugs. Nucleic Acids Res. 2017 Jan 4;45(D1):D353–61.

45. Salgado R, Denkert C, Demaria S, Sirtaine N, Klauschen F, Pruneri G, et al. The evaluation of tumor-infiltrating lymphocytes (TILS) in breast cancer: Recommendations by an International TILS Working Group 2014. Ann Oncol. 2015;26(2):259–71.

